# Amyloid Beta Peptides Inhibit Glucose Transport at the Blood-brain Barrier by Disrupting Insulin-Akt Pathway in Alzheimer’s Disease

**DOI:** 10.1101/2022.11.21.517280

**Authors:** Lushan Wang, Geoffry L. Curran, Paul H. Min, Ling Li, Val J. Lowe, Karunya K. Kandimalla

**Affiliations:** Department of Pharmaceutics and Brain Barriers Research Center, University of Minnesota, Minneapolis, Minnesota 55455, United States; Experimental and Clinical Pharmacology, College of Pharmacy, University of Minnesota, Minneapolis, Minnesota 55455, United States; Departments of Radiology, Mayo Clinic College of Medicine, Rochester, Minnesota, 55905, United States

**Author notes:** **Corresponding author:** Karunya K. Kandimalla, Department of Pharmaceutics, College of Pharmacy, University of Minnesota, Minneapolis, MN, 55455, USA.

## Abstract

Disruptions in glucose uptake and metabolism in the brain are implicated in metabolic disorders and Alzheimer’s disease (AD). Toxic soluble amyloid-beta (sAß) peptides accumulating in the brain and plasma of AD patients were suggested to promote blood-brain barrier (BBB) dysfunction, brain hypometabolism, and cognitive decline. Exposure to sAß peptides is reported to interfere with glucose metabolism in the brain parenchyma, although their effects on the BBB have not been fully characterized. Our data showed that the brain uptake of glucose surrogate, [^18^F]-fluorodeoxyglucose (^18^FDG), was reduced significantly in APP/PS1 transgenic mice (overproduce Aß) compared to wild-type (WT) mice. In addition, the influx rate of ^18^FDG was also decreased in both Aß40 and Aß42 pre-infused mice compared to control mice. Glucose is primarily delivered from blood into the brain via glucose transporter 1 (GLUT1). The confocal microscopy experiment showed that Aß40 and Aß42 peptides significantly decreased GLUT1 expression in polarized human cerebral microvascular endothelial cell (hCMEC/D3) monolayers. Insulin-AKT pathway has been observed to induce glucose uptake via regulating the expression of TXNIP, the only α-arrestin protein known to bind to thioredoxin. We found that Aß40 and Aß42 peptides decreased p-AKT and increased TXNIP expression in the hCMEC/D3 cell monolayers. MK2206, a kinase inhibitor of AKT, was used to confirm that inhibition of insulin/AKT pathway reduced GLUT1 expression in an insulin-independent manner in the hCMEC/D3 cell monolayers. These results suggest that inhibitory effects of sAß on GLUT1 expression are mediated by inhibition of the insulin/AKT pathway. The role of TXNIP on endothelial GLUT1 expression was investigated using resveratrol, which has been reported to downregulate TXNIP overexpression. Consistently, resveratrol treatment led to a significant increase in GLUT1 expression in the hCMEC/D3 cell monolayers. Furthermore, by co-incubation of resveratrol and sAß peptides in hCMEC/D3 cell monolayers, we found that resveratrol rectified the aberrant TXNIP expression caused by sAß peptides. Together, these findings provide novel evidence that toxic sAß peptide exposure inhibits glucose transport at the BBB by decreasing GLUT1 expression via the insulin/Akt/TXNIP axis.

## Introduction

Glucose is the main metabolic fuel for all mammalian cells. Despite being only ∼4% of total body weight, the human brain accounts for ∼ 25% of glucose consumption [1, 2]. Under normal conditions, glucose molecules are primarily delivered into the brain through facilitated diffusion by glucose transporter 1 (GLUT1) [3, 4]. The GLUT1 is expressed in the BBB endothelial cells and astrocytes but not in neurons [5, 6]. In BBB endothelial cells, a 55-kDa isoform of GLUT1 is expressed, whereas in astrocytes, a 45-kDa isoform is expressed [7].

Fluorodeoxyglucose (FDG), analogous to deoxyglucose, is transported into the brain at nearly the same rate as glucose, making it a suitable tracer to mimic glucose transport. As determined by [^18^F]-FDG positron emission tomography/computed tomography (PET/CT) imaging, cerebral hypometabolism has been shown in individuals with Alzheimer’s disease (AD) or an increased risk of AD [8-13]. The extent and rate of brain glucose uptake correlate with the level of GLUT1 expression in cerebral microvessels [14-16]. Reduced GLUT1 expression levels in cerebral microvessels were also found in transgenic mouse models of AD [17, 18] as well as in AD patients [19-22]. Studies in rodents indicate that reduced expression of GLUT1 in the endothelium initiates BBB breakdown and accelerates the progression of AD neuropathology [23].

In transgenic mouse models of AD with overexpression of amyloid-beta precursor protein (APP), higher amyloid-beta (Aß) levels, disrupt blood-brain barrier (BBB) integrity, causes BBB dysfunction, and reduces glucose transport [23]. However, the mechanisms by which Aß peptides impact glucose transport at the BBB are not well understood. Two major Aß isoforms, Aß40 and Aß42, predominate in the AD brain. Aß40 is the more vasculotropic isoform accumulating in the cerebral microvasculature, whereas Aß42 is more neurotoxic, amyloidogenic, and forms the nidus for amyloid plaque formation in the parenchyma of AD brains [24]. To explore relationships between Aß peptides and brain glucose uptake, we pre-infused the wild-type mice (B6SJLF1) with vehicle or Aß peptides and evaluated the corresponding plasma-to-brain influx rate of glucose using dynamic ^18^FDG-PET/CT imaging.

Previous studies have indicated that GLUT1 mRNA level is increased by the suppression of TXNIP in bovine oocytes and HepG3 cells [25, 26]. It was further reported that TXNIP binds to GLUT1 and facilitates its endocytosis through clathrin-coated pits (CCPs) [25]. Thus, TXNIP appears to be a negative regulator of GLUT1 and maintains glucose homeostasis. However, the impact of Aß peptides on TXNIP expression and underlying molecular mechanisms at the BBB endothelium are not well understood. Insulin-PI3K/Akt/mTOR pathway was reported to play a critical role in the re-localization of GLUT1 on cell membranes by downregulating the expression of TXNIP in non-small cell lung cancer cell lines [27]. Brain insulin resistance in AD patients is evidenced by insulin/AKT signaling disruptions in the brain [28]. Therefore, we hypothesized that Aß peptides could inhibit glucose transport at the BBB via insulin/Akt/TXNIP axis.

## Materials and Methods

### Materials

Aß40 and Aß42 peptides were procured from AAPPtec, LLC (Louisville, KY). Insulin (Novolin^®^ or Humulin^®^) was purchased from Eli Lily (Indianapolis, IN). ^18^Fluorodeoxyglucose (^18^FDG) was prepared in-house at the Mayo Clinic as per the clinical standards.

### Animals

Three-month-old female APP/PS1 transgenic mice overexpressing mutant forms of amyloid-beta precursor protein (APPswe) and presenilin-1 (PS1dE9) and wild-type (WT) mice, C57BL/6J or B6SJLF1, were purchased from the Jackson Laboratory (Bar Harbor, ME). They were housed in the Mayo Clinic animal care facility under standard conditions with access to food and water *ad libitum*. All studies were conducted in accordance with the National Institutes of Health guidelines for the care and use of laboratory animals and protocols approved by the Mayo Clinic Institutional Animal Care and Use Committee (Mayo IACUC #A00006176-21). Prior to the experiments, the mice were visually inspected to be in the diestrus phase of the estrous cycle [2], during which hormonal fluctuations are expected to be minimal [29].

### Determination of ^18^FDG brain distribution in APP/PS1 mice via dynamic PET/CT imaging

Each mouse was fasted for 12 hours before the experiment. APP/PS1 or WT (C57BL/6J) mice were anesthetized using a mixture of isoflurane (1.5 %) and oxygen (4 L/min). A bolus of ^18^FDG (500 μCi/200 µL) was injected into the femoral vein, and the mouse was imaged in the PET/CT scanner (Siemens Inveon® Multi-Modality System, Knoxville, TN) from 0 to 40 min. The average pixel values multiplied by volume-of-interest volume (AVR*VOL) were calculated using PMOD software (PMOD Technologies LLC, Fällanden, Switzerland), and it represents total ^18^FDG activity in the volume-of-interest (VOI). The AVR*VOL value in the mouse brain at the end of 40 min was used to compare cerebral ^18^FDG uptake difference between APP/PS1 mice and their WT counterparts.

### Determination of sAß effect on ^18^FDG plasma pharmacokinetics and brain influx rate

Aß peptides were prepared as described in our previous publication [30]. The impact of sAß peptides on glucose uptake in the brain was determined by pre-infusing 150 µL of vehicle (0.17% DMSO in PBS) containing 250 μg of Aß40 or Aß42 into the left internal carotid artery of WT mice (B6SJLF1) at the rate of 10 μL/min. Fifteen minutes after the end of infusion, a bolus of ^18^FDG (500 μCi/200 µL) was injected via the femoral vein. The mouse was imaged by dynamic PET/CT from 0 to 30 min followed by a 5 min CT scan to locate regions of interest. The blood (20 μL) was simultaneously sampled from the femoral artery from 0.25 to 30 min. The plasma pharmacokinetics and the brain influx rate were determined as detailed below.

### Plasma pharmacokinetics of ^18^FDG

The blood was separated as red blood cells and plasma, and ^18^F activity was assayed using a gamma counter (***Figure 2 A***). The ^18^FDG plasma time-activity (*C*_*p*_(*t*)) are corrected for the ^18^F physical decay [31-35] as follows:

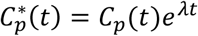

where λ (minute^-1^) is the ^18^F physical decay constant (*λ* = ln 2/110) and t is expressed in minutes.

The ^18^FDG plasma concentration versus time profile was fitted with the following bi-exponential equation via Phoenix^®^ WinNonlin 6.4 (Certara, St. Louis, MO):

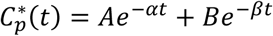

where 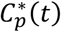 is the concentration (µCi/mL) in plasma at time t (min), A and B are the intercepts of the distribution and elimination phases, respectively, and α and ß are the distribution and elimination micro-rate constants (minute^-1^), respectively. The secondary plasma pharmacokinetics parameters, including Cmax, area under the curve (AUC), and clearance (CL), are calculated by the software.

### Brain influx rate of ^18^FDG

The ^18^FDG brain time-activity (*Amount*_*brain*_(*t*)) curves are first corrected for the ^18^F physical decay as follows:

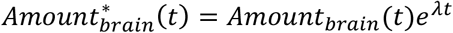

where λ (minute^-1^) is the ^18^F physical decay constant (*λ* = ln 2/110) and t is expressed in minutes.

The plasma-to-brain influx rate of ^18^FDG in WT mice with or without Aß pre-infusion was determined by Gjedde-Patlak graphical analysis [36, 37], which is constructed by plotting

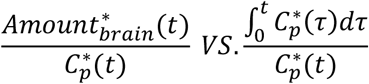

where 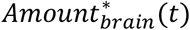 is ^18^F activity (µCi) in the brain region of interest (ROI); 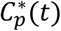 is plasma concentration (µCi/mL) at time t; 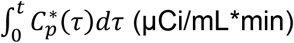 is area under the plasma concentration (µCi/mL) versus time (0-t min) profile determined by the trapezoidal method. The slope obtained from linear regression of the plot represents plasma-to-brain influx rate, K_i_ (mL/min).

### Cell culture

Polarized human cerebral microvascular endothelial cell (hCMEC/D3) monolayers were used as a BBB model. These cell lines were a kind gift from P-O Couraud (Institut Cochin, France) and were cultured as previously described [38].

### Western blotting

To test the impact of sAß exposure and MK2206 (Akt inhibitor) on Akt phosphorylation, western blotting was performed as described in our previous publication [39]. The concentration of Aß40 and Aß42 in the culture medium was quantified by ELISA (Thermo Scientific, Waltham, MA). Briefly, hCMEC/D3 cell monolayers were cultured on 6-well plates. Upon reaching confluency, the cells were incubated with the low serum medium containing 1% v/v FBS overnight. Next day, the cells were pre-incubated with Dulbecco’s modified Eagle’s medium (DMEM), 1.25 µg/mL Aß40 or Aß42 in DMEM for 1 hour, or 750 nM MK2206 in DMEM for 4 hours at 37 ^o^C. Then they were incubated with 100 nM insulin for another 20 min at 37°C. To investigate the impact of sAß exposure or resveratrol on TXNIP expression, hCMEC/D3 cell monolayers were incubated with Dulbecco’s modified Eagle’s medium (DMEM), 1.25 µg/mL Aß40 or Aß42 in DMEM for 1 hour, or 160 µM resveratrol for 12 hours in the low-serum medium at 37°C. After the incubation, the cells were washed three times with PBS and lysed in RIPA buffer containing protease and phosphatase inhibitors (Sigma-Aldrich, St. Louis, MO). Total protein concentrations in the lysates were determined by bicinchoninic acid (BCA) assay (Pierce, Waltham, MA). Lysates were loaded onto 4-12% Criterion XT precast gels and resolved by SDS-PAGE under reducing conditions (Bio-Rad Laboratories, Hercules, CA). The proteins were then electro-blotted onto a 0.45 µm nitrocellulose membrane. The membranes were blocked with 5% non-fat dry milk protein (Bio-Rad Laboratories), incubated overnight at 4°C with primary antibodies against Vinculin, Akt, p-Akt (S473), or TXNIP (1:500 for TXNIP; 1:1000 for other antibodies; Cell Signaling Technology, Danvers, MA). Then the membranes were incubated with fluorescent-conjugated secondary antibody (1:2000) for 1 hour at room temperature. Immuno-reactive bands thus formed were imaged by LI-COR (Odyssey CLx, Lincoln, NE), and the band intensities were quantified by densitometry using Image Studio (Image Studio Lite Ver 5.2, Lincoln, NE).

### Immunocytochemistry

The hCMEC/D3 cell monolayers were cultured on 35-mm coverslip bottom dishes as described previously [38]. Upon reaching confluency, the cell monolayers were incubated with or without Aß40 or Aß42 for 1 hour or 160 µM resveratrol for 12 hours at 37 °C. To test the impact of insulin-Akt pathway on GLUT1 expression at the BBB, the hCMEC/D3 cell monolayers cultured on dishes were incubated with or without 750 nM MK2206 for 4 hours, followed by 100 nM insulin stimulation for another 20 min. After incubation, the cell monolayers were washed with prechilled PBS, fixed with 100% prechilled methanol for 5 min at room temperature, and blocked with 10% goat serum (Sigma-Aldrich, St. Louis, MO) for 30 min. The cell monolayers were incubated overnight with GLUT1 primary antibody (1:100), washed three times using DPBS, and incubated with goat anti-rabbit IgG secondary antibody (1:250) for 1 hour in the dark at room temperature. After rinsing thoroughly with DPBS, dishes were mounted with ProLong Diamond mounting medium containing 4’,6-diamidino-2-phenylindole (DAPI) (Invitrogen, Waltham, MA), and imaged using a Zeiss LSM 780 laser confocal microscope equipped with a C-Apochromat 40x/1.2W objective.

### Statistical analysis

For comparisons between two groups, an unpaired, two-tailed student’s t-test was used. One-way ANOVA with Bonferroni post hoc was used for comparing groups of three or more. All statistics were performed using GraphPad Prism (GraphPad Software; La Jolla, CA).

## Results

### Significant reduction of ^18^FDG uptake in APP/PS1 transgenic mice

The ^18^FDG uptake in the brain of young APP/PS1 and WT mice (3-month-old) was monitored by 40 minutes of dynamic PET/CT imaging. At the end of imaging, the cerebral uptake of ^18^FDG (AVR*VOL) in APP/PS1 transgenic mice was found significantly decreased by ∼ 40 % compared to their WT counterparts (***Figure 1***).

**Figure 1:**
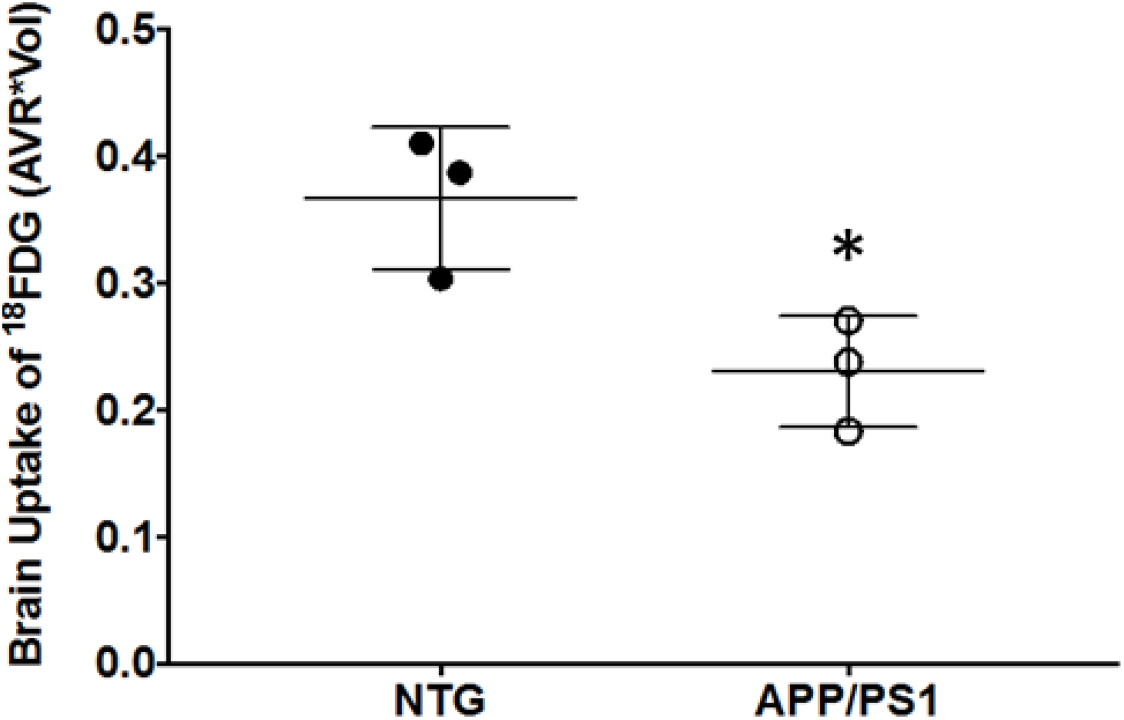
Brain uptake of ^18^FDG in non-transgenic(NTG) vs. APP/PS1 transgenic mice. An intravenous bolus (500 µCi) injection of ^18^FDG was administered via femoral vein, and the mice were monitored by dynamic PET/CT imaging for 40 min. Bar chart of total ^18^FDG radioactivity in the VOI (AVR*VOL, R.U.) of NTG vs. APP/PS1 mice at the end of the imaging. Data represent mean ± SD (n=3). Unpaired student’s t-test (*p<0.05).

### Significant reduction of ^18^FDG uptake in sAß pre-infused mice

To further confirm the impact of sAß exposure on ^18^FDG uptake, WT mice were with PBS with or without sAß peptides. After intravenous bolus injection, the plasma concentration of ^18^FDG in 3-month-old WT mice declined in a bi-exponential fashion over the first 30 minutes with and without Aß pre-infusion, and no significant differences in the plasma pharmacokinetics were observed among the groups (***Figure 2 B***). As shown in ***Figure 2 C***, the ratio of total brain amount to the plasma concentration at the respective times versus the ratio of the plasma concentration-time integral to *C*^*∗*^(*t*) yielded a linear curve, indicating unidirectional transfer from plasma to the brain within the certain time slot. The plasma-to-brain influx rate (Ki) of ^18^FDG was found to be lower in both Aß40 and Aß42 pre-infused mice compared to the control mice. The Ki of ^18^FDG was 0.043±0.009 mL/min in PBS pre-infused mice (n=6), decreased to 0.012±0.003 mL/min in Aß40 pre-infused mice (n=3), and 0.007±0.006 mL/min in Aß42 pre-infused mice (n=3) (***Figure 2 D***).

**Figure 2:**
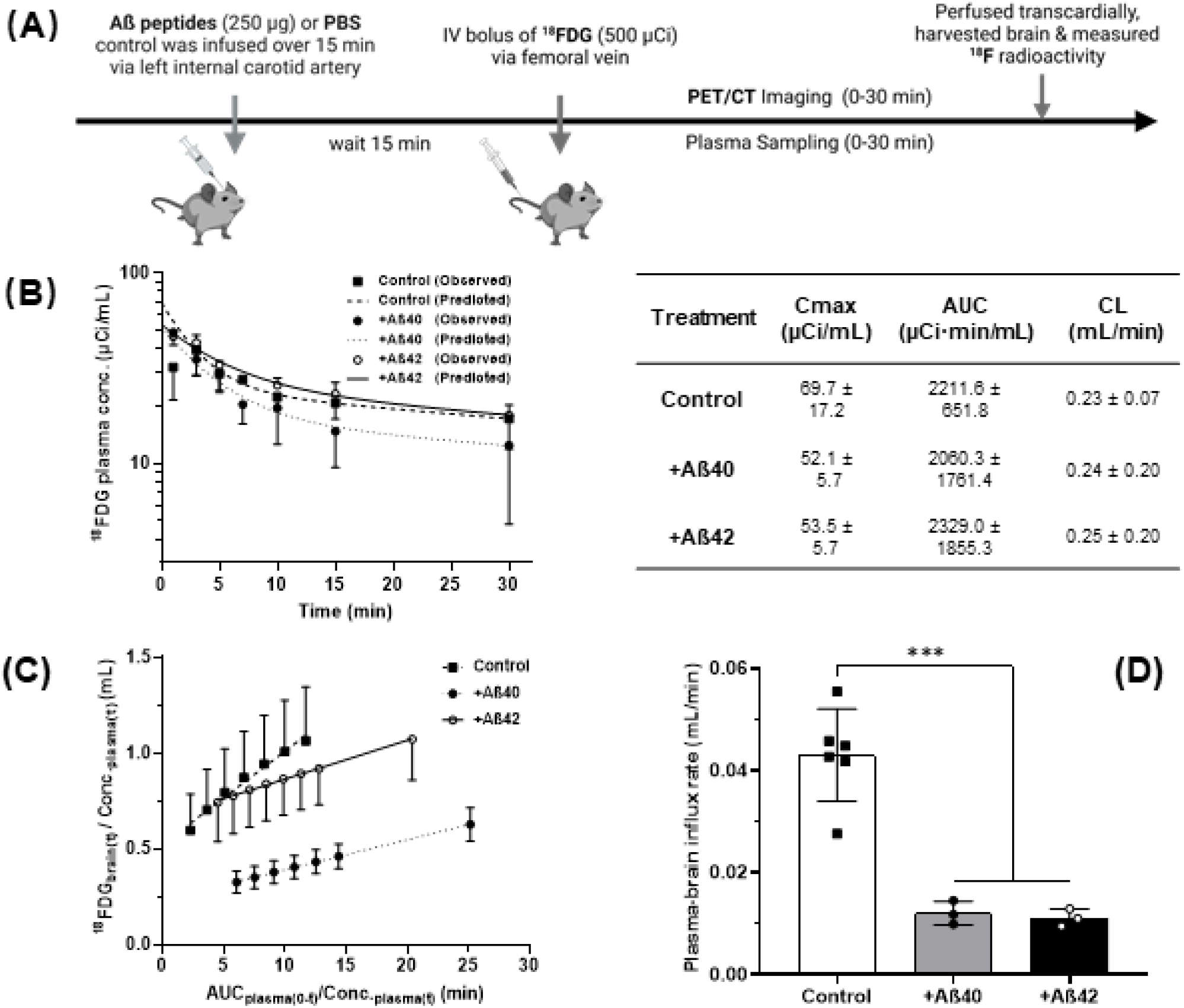
Effects of Aß peptides on plasma-to-brain influx rate of ^18^FDG in mice. **(A)** Experimental scheme. Mice were pre-infused with Aß40 or Aß42 peptides (250 µg) or PBS control via left internal carotid artery over 15 min, followed by a bolus injection of ^18^FDG (500 µCi/200 µL) into femoral vein. Tracer accumulation in the brain was monitored by dynamic PET/CT imaging from 0 to 30 min, and plasma was sampled simultaneously from femoral artery. After obtaining the final sample, the mice were transcardially perfused with PBS, and the ^18^FDG radioactivity in plasma and dissected brain were assayed in the gamma counter. **(B)** Plasma concentration versus time profile of ^18^FDG in mice pre-infused with Aß40, Aß42, or PBS control. Shown are the predicted curves laid over the observed values. Plasma pharmacokinetic parameters were not significantly different among the three groups of mice. **(C)** The plasma-to-brain influx rate (Ki) of ^18^FDG was estimated from the slope of Gjedde-Patlak plot. **(D)** Comparison of plasma-to-brain influx rate (Ki; mL/min) with and without Aß exposure. Data represent mean ± SD (n=6 for control group, n=3 for Aß treated groups). One-way ANOVA followed by Bonferroni’s multiple comparisons tests (*** p<0.005).

### Aß40 and Aß42 reduced GLUT1 expression in polarized hCMEC/D3 cell monolayers

The GLUT1 expression in polarized hCMEC/D3 cell monolayers was investigated to determine the effect of sAß exposure. The confocal micrographs demonstrated lower AF647-GLUT1 fluorescent intensity in Aß40 (8 × 10^5^ ± 2.5 × 10^5^ a.u.) and Aß42 pretreated cell monolayers (6.9 × 10^5^ ± 1.1 × 10^5^ a.u.) compared to control cell monolayers (13.2 × 10^5^ ± 5.2 × 10^5^ a.u.) (***Figure 3 A***).

**Figure 3.**
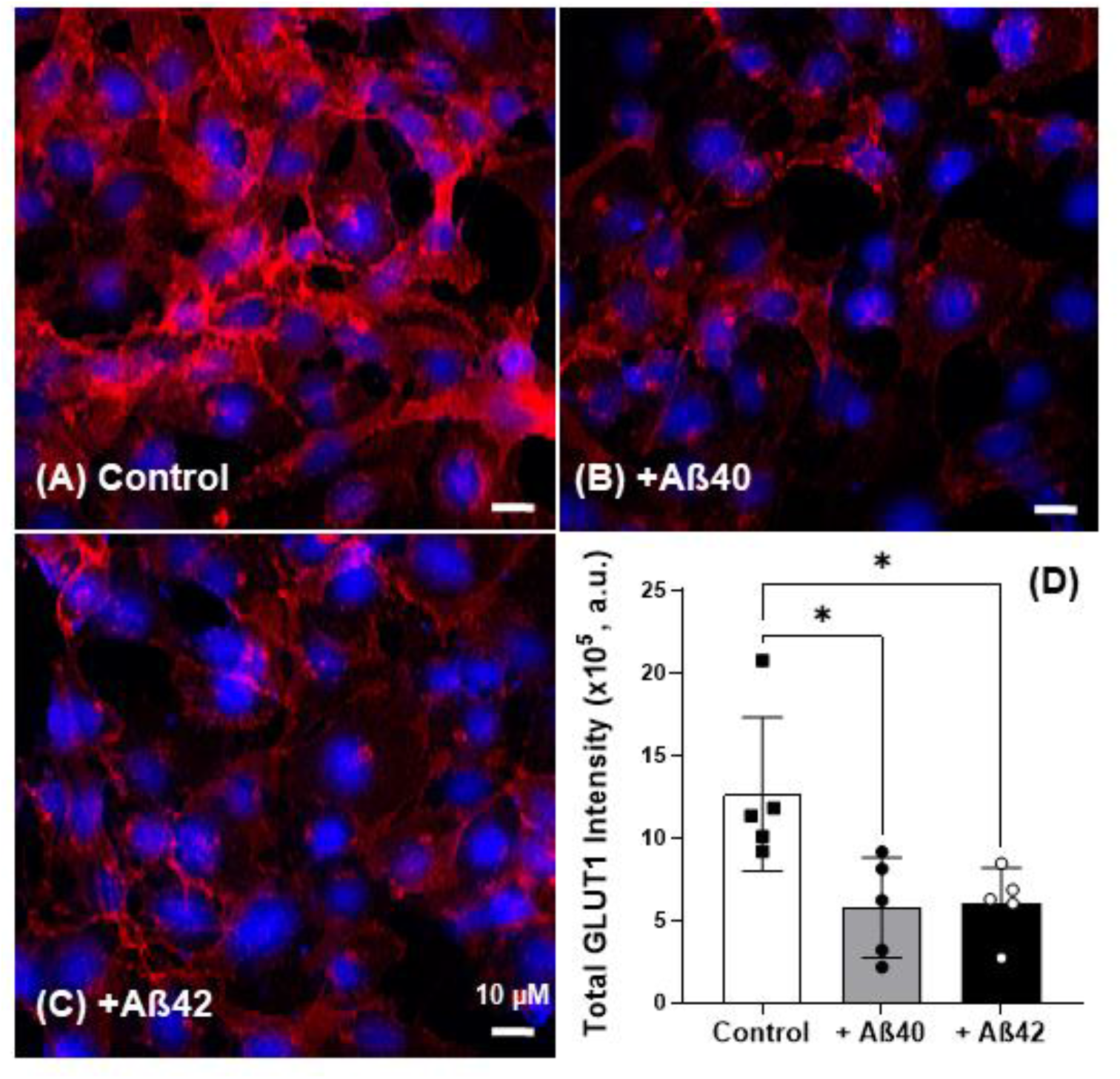
Both Aß40 and Aß42 reduce GLUT1 expression in hCMEC/D3 cell monolayers. **(A-C)** Representative laser confocal micrographs of hCMEC/D3 cell monolayers pretreated with **(B)** Aß40 or **(C)** Aß42 show reduced intracellular AF647-GLUT1 expression when compared with cells treated with **(A)** control. **(D)** Quantification of total AF647-GLUT1 fluorescence intensity shows that both Aß40 and Aß42 exposure significantly lower the GLUT1 expression in hCMEC/D3 cell monolayers compared to the control. Red= AF647-GLUT1 and blue = DAPI-stained nuclei; Scale bar, 10 µm. Data represent mean ± SD (n=4). One-way ANOVA with Bonferroni post hoc (* p<0.05).

### Aß40 and Aß42 disrupted Insulin-Akt pathway and upregulated TXNIP expression in polarized hCMEC/D3 cell monolayers

Our previous publication has shown that Aß peptides disrupt insulin signaling in the cerebral vascular endothelial cells [40]. Consistently, in the current study, we have shown that Aß40 and Aß42 significantly decreased phosphorylation of Akt by 2 to 3-fold in hCMEC/D3 cell monolayers (***Figure 4***). Moreover, Aß40 and Aß42 significantly increased the expression of TXNIP (***Figure 5 A-C***), which is a negative regulator of GLUT1 plasma membrane localization [25, 41]. These results were confirmed using confocal microscopy, where the normalized fluorescent intensity (total fluorescent intensity/number of cells) of AF647-TXNIP in hCMEC/D3 cell monolayers significantly increased after sAß exposure compared to control monolayers (***Figure 8)***.

**Figure 4.**
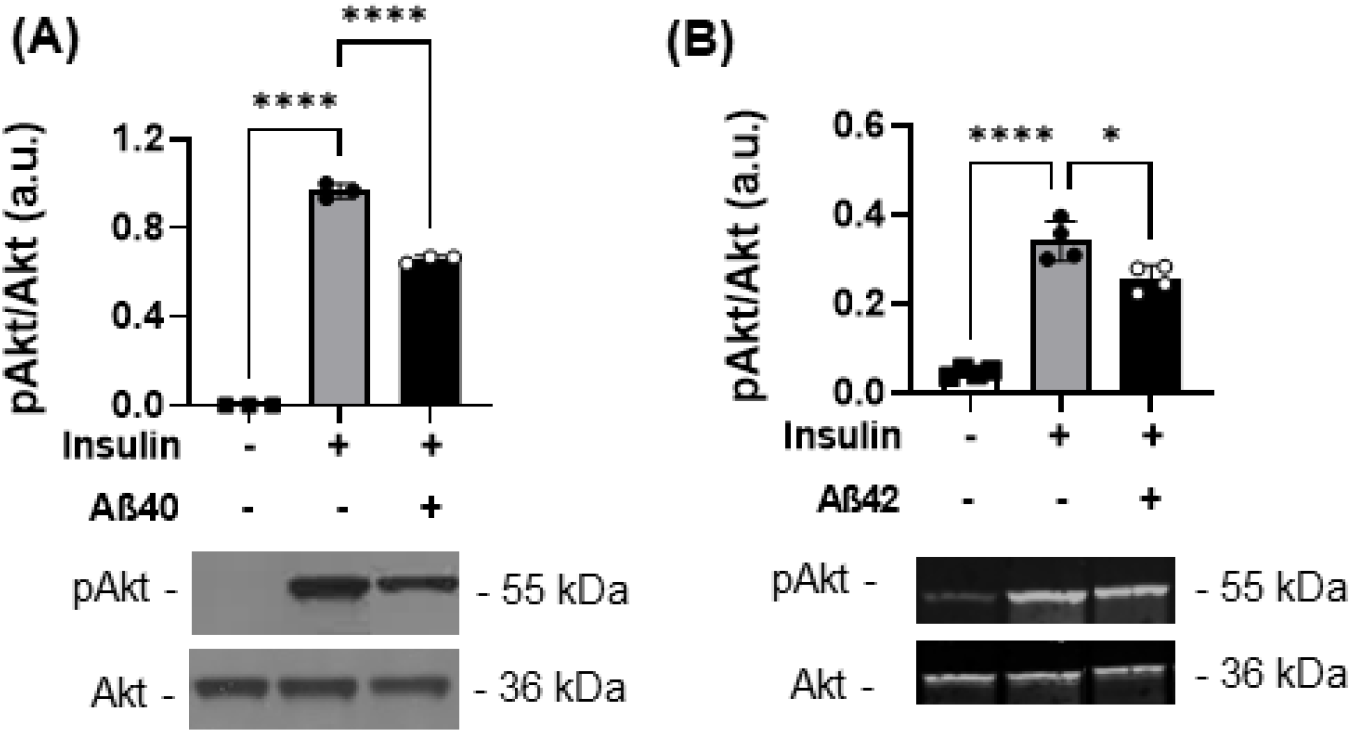
Both (A) Aß40 and (B) Aß42 inhibit phosphorylation of AKT in hCMEC/D3 cell monolayers. Western blots were performed to assess the expression and phosphorylation of AKT in hCMEC/D3 cell monolayers following 1-hour treatment with or without Aß exposure (1.25 µg/mL) and stimulation with insulin (100 nM) for another 20 min. Figure is shown as bar charts (*top*) and its corresponding representative immunoblots (*bottom*) of p-AKT/AKT. Data represent mean ± SD (n=3 for Aß40, n=4 for Aß42). One-way ANOVA with Bonferroni post hoc (* p<0.05; **** p<0.001).

**Figure 5.**
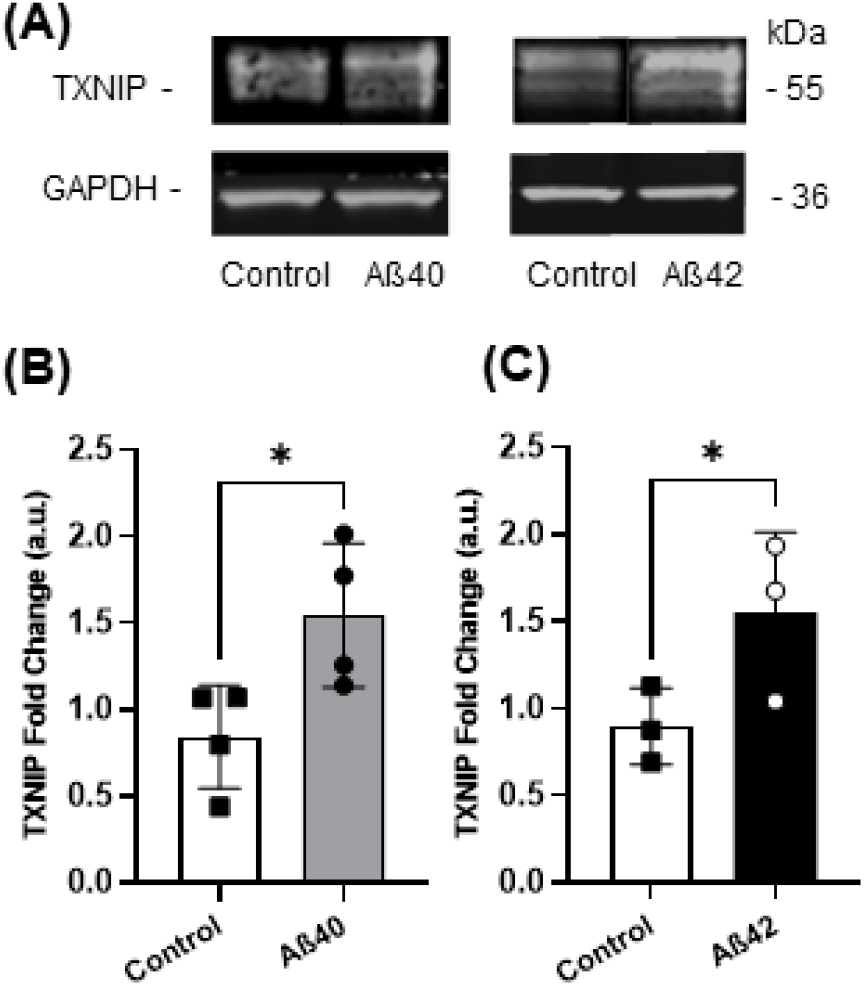
Both Aß40 and 42 increase TXNIP expression in hCMEC/D3 cell monolayers. (**A-C**) Western blots were performed to assess the expression of TXNIP in hCMEC/D3 cell monolayers following 1-hour treatment with control, Aß40 (1.25 µg/mL; **B**) or Aß42 (0.625 µg/mL; **C**). Shown are (**A**) representative immunoblots and (**B-C**) bar charts of quantification for TXNIP fold changes. Data is shown as mean ± SD (n=3); Unpaired student t-test (*p<0.05).

### Akt inhibitor MK2206 reduced GLUT1 expression in polarized hCMEC/D3 cell monolayers

MK2206 is known to disrupt Akt phosphorylation [42] and is widely used as an Akt inhibitor. To investigate the role of Akt activation on GLUT1 expression in hCMEC/D3 cell monolayers, we performed immunofluorescence study in the hCMEC/D3 cell monolayers treated with or without MK2206 followed by insulin stimulation. The GLUT1 expression of cells after insulin stimulation (1.36 ± 0.15; n=4) was not significantly altered compared to untreated monolayers (2.02 ± 0.42; n=4) (***Figure 6 A&C***). However, MK2206 reduced the expression of GLUT1 with or without insulin stimulation (***Figure 6 B&D***).

**Figure 6.**
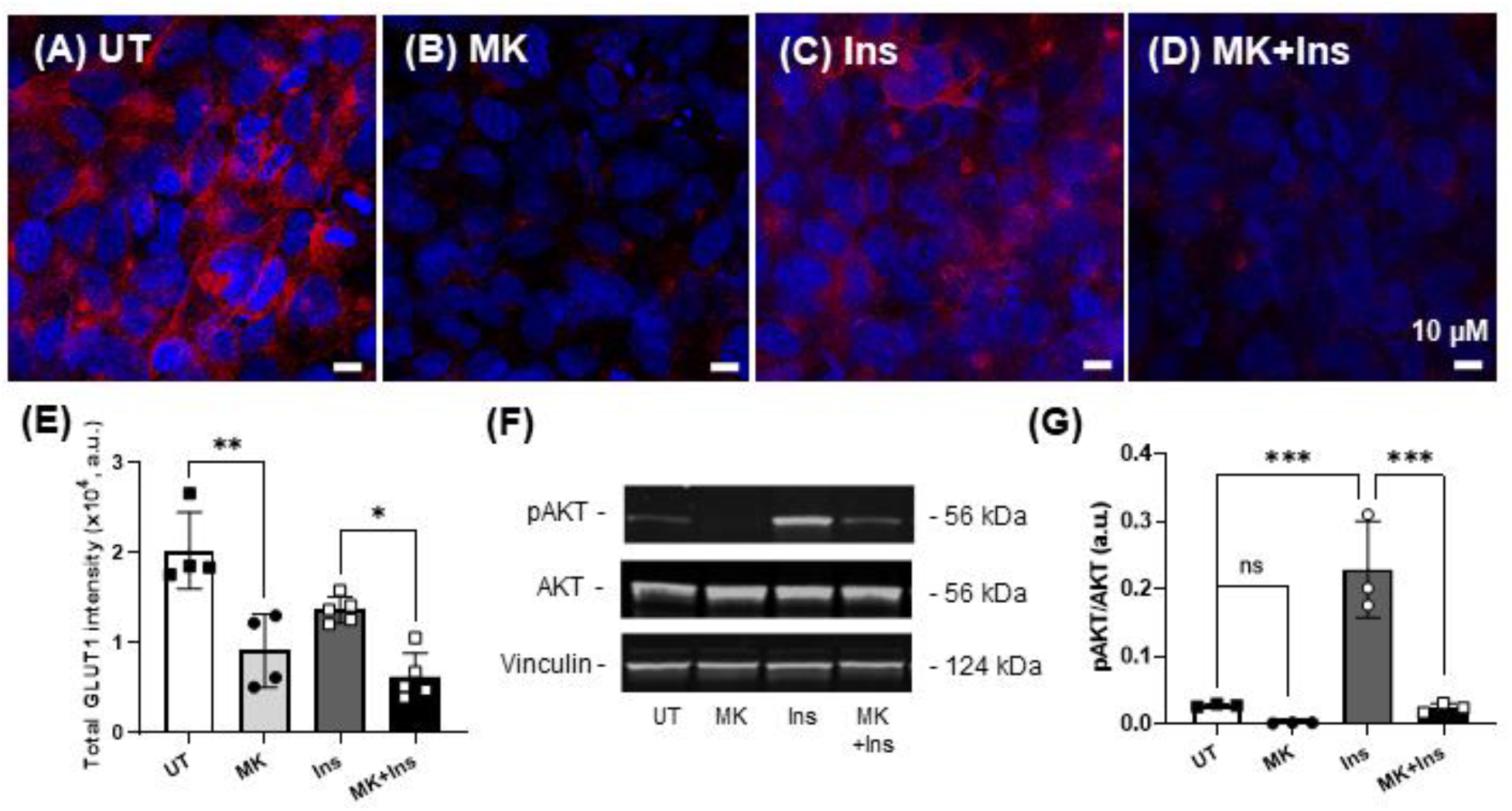
AKT inhibitor (MK2206) reduces GLUT1 expression in hCMEC/D3 cell monolayers. **(A-D)** Representative laser confocal micrographs of intracellular expression of GLUT1 in hCMEC/D3 cell monolayers following 4-hour treatment with or without AKT inhibitor MK2206 (750 nM) and stimulation with insulin (100 nM) for another 20 min. Red= AF647-GLUT1 and blue = DAPI-stained nuclei; Scale bar = 10 µm. **(E)** The bar chart represents the quantification of total AF647-GLUT1 fluorescence intensity. Data is shown as mean ± SD (n=4 for UT and MK groups; n=5 for Ins and MK+Ins groups). **(F)** Shown are representative immunoblots of pAKT, AKT, and Vinculin in hCMEC/D3 cell monolayers treated with AKT inhibitor MK2206 (750 nM) and stimulated with insulin (100 nM). **(G)** The bar chart represents the quantification of pAKT/AKT. Data represent mean ± SD (n=3). One-way ANOVA with Bonferroni post hoc (**<0.01; *** p<0.005). Abbreviations: UT: untreated; MK: MK2206; Ins: insulin.

### Role of TXNIP on GLUT1 expression in polarized hCMEC/D3 cell monolayers

To confirm the role of TXNIP on GLUT1 expression in the hCMEC/D3 cell monolayers, we used a natural polyphenol, resveratrol, to inhibit TXNIP expression. As shown in **Figure 7 D & E**, resveratrol decreased TXNIP expression by 70% in hCMEC/D3 cell monolayers as determined using western blots. In contrast, the fluorescence intensity of AF647-GLUT1 increased by 50% in the resveratrol pretreated monolayers (***Figure 7 B&C***) compared to control monolayers (***Figure 7 A&C***).

**Figure 7:**
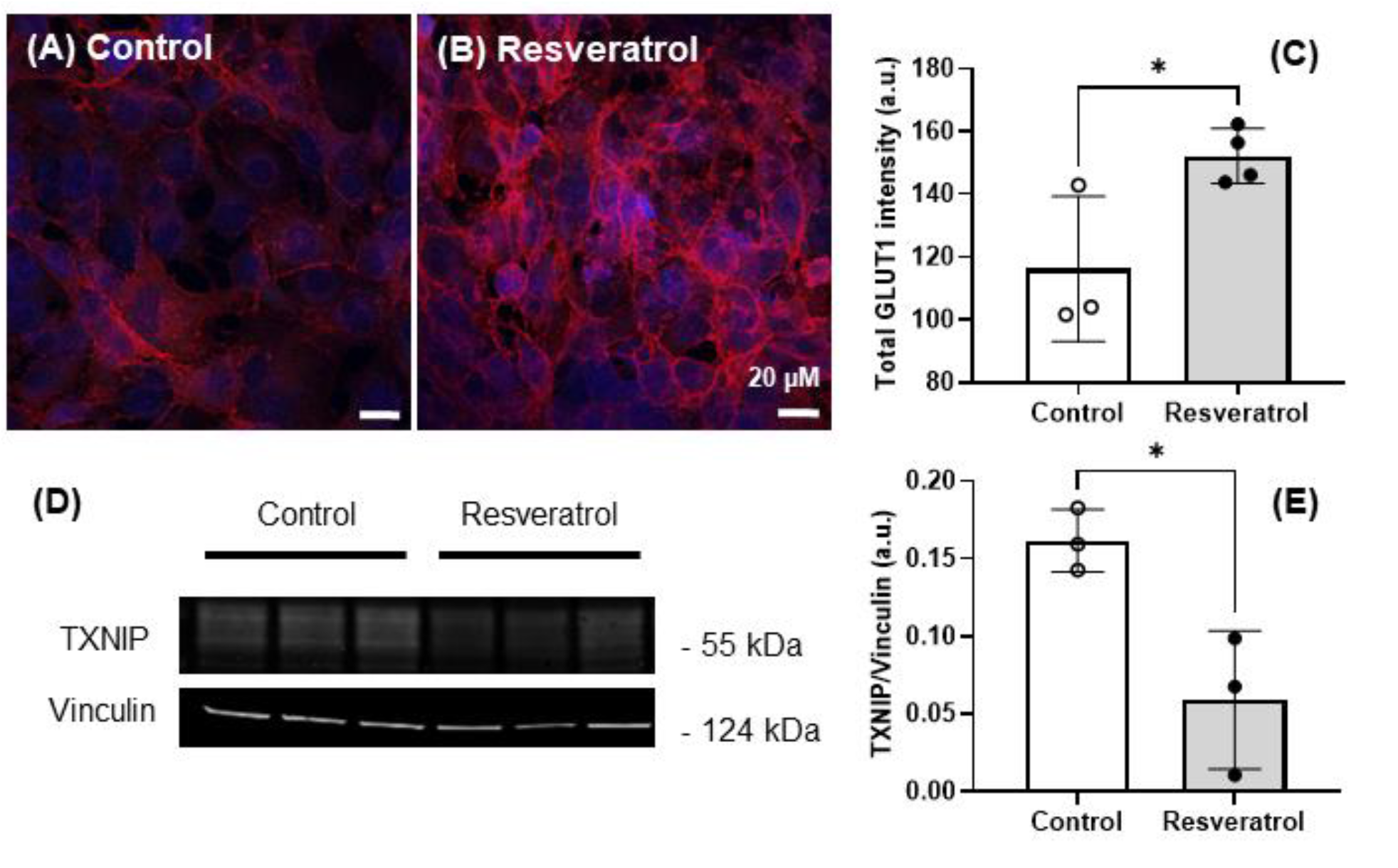
TXNIP is a negative regulator of GLUT1 expression in hCMEC/D3 cell monolayers. The hCMEC/D3 cell monolayers were treated for 12 hours with or without 160 µM of resveratrol in a low serum medium. Representative laser confocal micrographs of AF647-GLUT1 expression in hCMEC/D3 cell monolayers treated with **(A)** control or **(B)** resveratrol. **(C)** Bar chart of total AF647-GLUT1 fluorescence intensity. Data represent mean ± SD (n=3). **(D)** Western blots and **(E)** densitometric analysis of TXNIP and Vinculin in control and resveratrol-treated cell monolayers. Red = AF647-GLUT1 and blue = DAPI; Scale bar = 20 µm. Data represent mean ± SD (n = 3). Unpaired student t-test (*p<0.05).

### Resveratrol normalized TXNIP expression stimulated by sAß exposure in polarized hCMEC/D3 cell monolayers

We further tested if resveratrol affects TXNIP expression induced by sAß exposure in hCMEC/D3 cell monolayers using immunofluorescence studies. Resveratrol treatment alone decreased TXNIP by 50% compared to control monolayers. More importantly, the co-incubation of resveratrol and sAß peptides significantly reduced the TXNIP expression compared to sAß alone treated monolayers (***Figure 8***).

**Figure 8:**
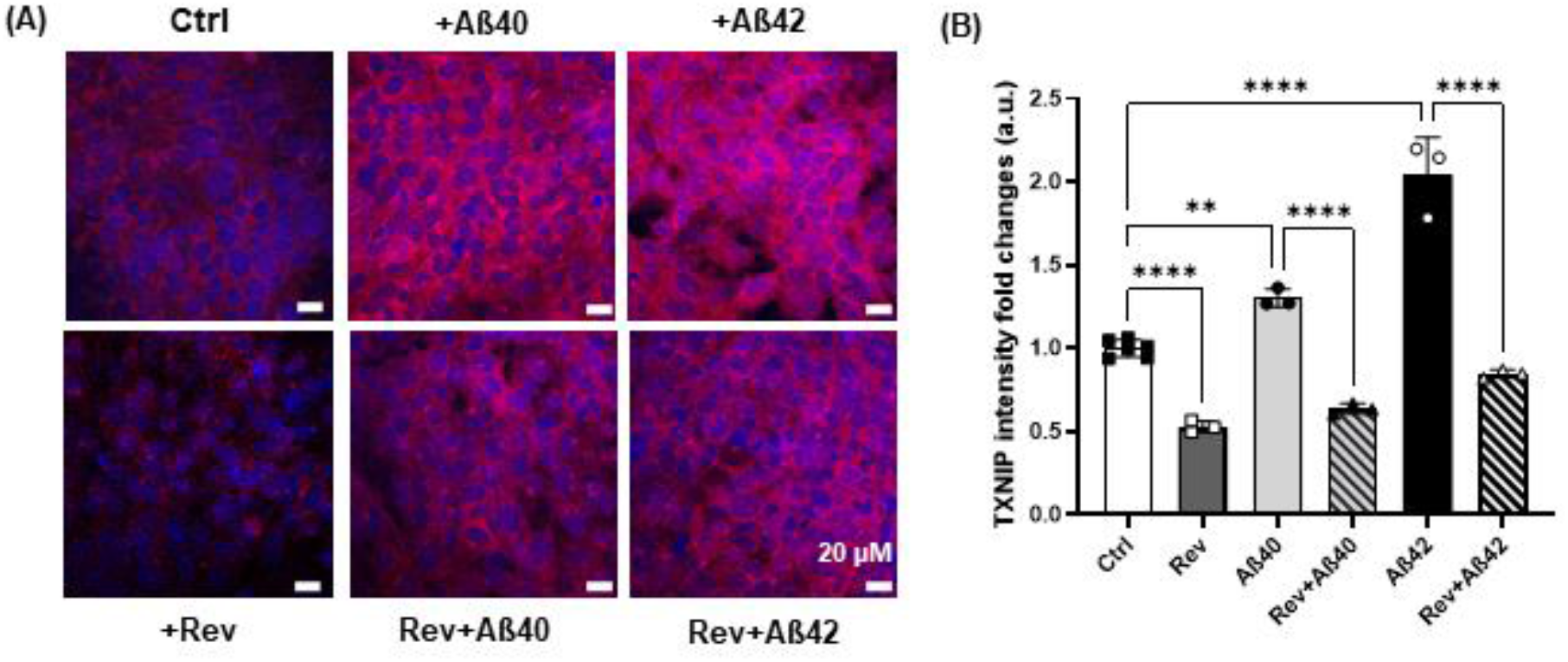
Resveratrol restore TXNIP expression induced by sAß peptides in hCMEC/D3 cell monolayers. **(A)** The hCMEC/D3 cell monolayers were treated for 12 hours with or without 160 µM of resveratrol in low serum medium following sAß treatment (Aß40:1.25 µg/mL; Aß42: 0.625 µg/mL) for another 1 hour. Representative laser confocal micrographs of TXNIP expression in hCMEC/D3 cell monolayers with various treatments. (**B**) The bar chart represents fold changes in AF647-TXNIP fluorescence intensity among the groups. Red = AF647-GLUT1 and blue = DAPI; Scale bar = 20 µm. Data represent mean ± SD (n=6 for Control group and n=3 for other groups). One-way ANOVA with Bonferroni post hoc (**** p<0.001).

## Discussions

Reduced glucose metabolism in hippocampus and cortical regions has been observed in individuals who are genetically at risk for AD [9, 10]. Transgenic AD mouse models that overexpress APP have also been shown to display altered glucose uptake and metabolism in the brain [8, 43-48]. The glucose uptake was shown to be either decreased or increased depending on the age/stage of the disease and the model. Our in vivo study indicates that 3-month-old APP/PS1 mice exhibit deficits in the extent of ^18^FDG uptake to the brain **(*Figure 1*)**. Notably, at this young age, the APP/PS1 mouse brain is devoid of Aß plaques as the amyloid deposition does not manifest until 6 months of age in these mice [49]. The fact that deficits of glucose uptake occur in young, plaque-free APP/PS1 mice suggests that soluble Aß (sAß) peptides are most likely responsible for the reduction of ^18^FDG uptake in AD mice. To avoid confounding effects of APP and presenilin mutations on glucose transport and metabolism, we pre-infused 3-month-old wild-type (WT) mice with sAß peptides to determine their impact on the brain glucose transport. The influx rate of ^18^FDG was estimated by non-invasive dynamic ^18^FDG-PET/CT imaging followed by Patlak graphical analysis, a widely used kinetic analysis describing irreversible tissue accumulation of the tracer [36, 37]. As shown in ***Figure 2*D**, both Aß40 and Aß42 significantly decreased plasma-to-brain influx rate of ^18^FDG in WT mice. Most of the glucose transport from blood to the brain is handled by the blood-brain barrier (BBB), and to a minor extent by the blood– cerebrospinal fluid (CSF) barrier [47]. The exposure of sAß peptides have been linked to impairment of glucose uptake, other membrane transport processes, and signaling systems in cultured rat hippocampal and cortical neurons [50]. Thus, it is likely that sAß peptides may disrupt BBB glucose transport into the brain.

Glucose transporter 1 (GLUT1), expressed on the BBB, is the primary transporter delivering glucose from blood into the brain. It has been well documented that the early reduction in glucose transport found in AD patients is associated with diminished BBB GLUT1 expression [23, 51, 52]. Recent study also suggests that Aß can induce irregularity in the translocation of GLUT1 to the plasma membrane in the brain parenchymal cells [17]. Interestingly, cerebral hypoglycemia was evident in younger APP/PS1, even before the amyloid accumulation is evident, thereby, suggesting that sAß peptides disrupt GLUT1 expression in these mice. Indeed, our confocal microscopy experiments confirmed that sAß peptides significantly decreased GLUT1 expression in hCMEC/D3 cell monolayers (***Figure 3***), which may contribute to reduction in the rate and extent of glucose uptake into the brain.

However, the mechanisms by which sAß disrupts endothelial GLUT1 expression is not well understood. Our previous study [40] showed that sAß peptides impair insulin signaling in the BBB endothelial cells as well as in 3xTg-AD mice. Of the two major arms of insulin signaling pathway, PI3K/Akt and MAPK/Erk, the PI3K/Akt pathway has canonical functions in glucose metabolism. We further confirmed that sAß peptides disrupt PI3K/Akt arm by decreasing the phosphorylation of Akt (***Figure 4***). Using MK2206, an allosteric Akt inhibitor, to test the role of Akt activation on GLUT1 endothelial membrane expression. As shown in ***Figure 6***, MK2206, an allosteric Akt inhibitor, substantially decreased endothelial GLUT1 expression in an insulin-independent manner. Together, these findings indicate that the inhibitory effects of sAß exposure on insulin-Akt pathway leads to diminished GLUT1 expression on the BBB endothelium, which contributes to decreased glucose uptake into the brain.

A growing body of literature has revealed that inhibition of the PI3K/Akt/mTOR pathway effectively suppresses GLUT1 membrane localization and leads to accumulation in the cytoplasm of lung cancer cell lines [53-55]. In addition, TXNIP protein, which functions as an adaptor for endocytosis of the GLUT1, was shown to negatively regulate glucose uptake in human skeletal muscle cells, adipocytes, hepatocytes, and HepG2 cells [25, 56]. Therefore, we tested the impact of sAß exposure on TXNIP expression in hCMEC/D3 cell monolayers and found that sAß exposure increased TXNIP expression (***Figure 5)***. This observation is consistent with previous findings that TXNIP protein, as well as its mRNA levels, were significantly elevated in human AD brains with Aß plaques compared with non-dementia control brains [57].

Finally, the role of TXNIP on endothelial GLUT1 expression was investigated using resveratrol. Resveratrol is a plant-derived natural polyphenol with various beneficial effects, including neuroprotection and improved metabolic response [58]. It has been reported that resveratrol can inhibit TXNIP expression by inducing phosphorylation of AMPK [59-61]. This observation was verified by western blots conducted in hCMEC/D3 cell monolayers (***Figure 7***), in which endothelial GLUT1 expression was induced through inhibiting TXNIP expression by resveratrol. This is in line with the recent finding that resveratrol significantly decreases TXNIP mRNA and protein nuclear expression and helps restore several aging impairments [60]. We further tested and found that resveratrol can rectify the reduction in TXNIP expression caused by sAß exposure (***Figure 8***). Therefore, TXNIP is a potential therapeutic target for AD, and resveratrol may help restore metabolic regulation in AD brain.

## Conclusions

Current understanding of the pathological association between metabolic disorders and sAß exposure is majorly limited to epidemiological findings. To date, the related pathological mechanisms have not been extensively explored in AD. The current work clarified key cellular/molecular mechanisms by which sAß disrupts brain glucose transport via the BBB (***Figure 9***). Our studies demonstrate that reduction of endothelial GLUT1 expression is responsible for reduced glucose transport observed in both AD transgenic and sAß pre-infused mice. Moreover, sAß peptides induce TXNIP expression by suppressing insulin-Akt pathway, lead to diminished BBB GLUT1 expression, and eventually reduce glucose delivery to the brain.

**Figure 9:**
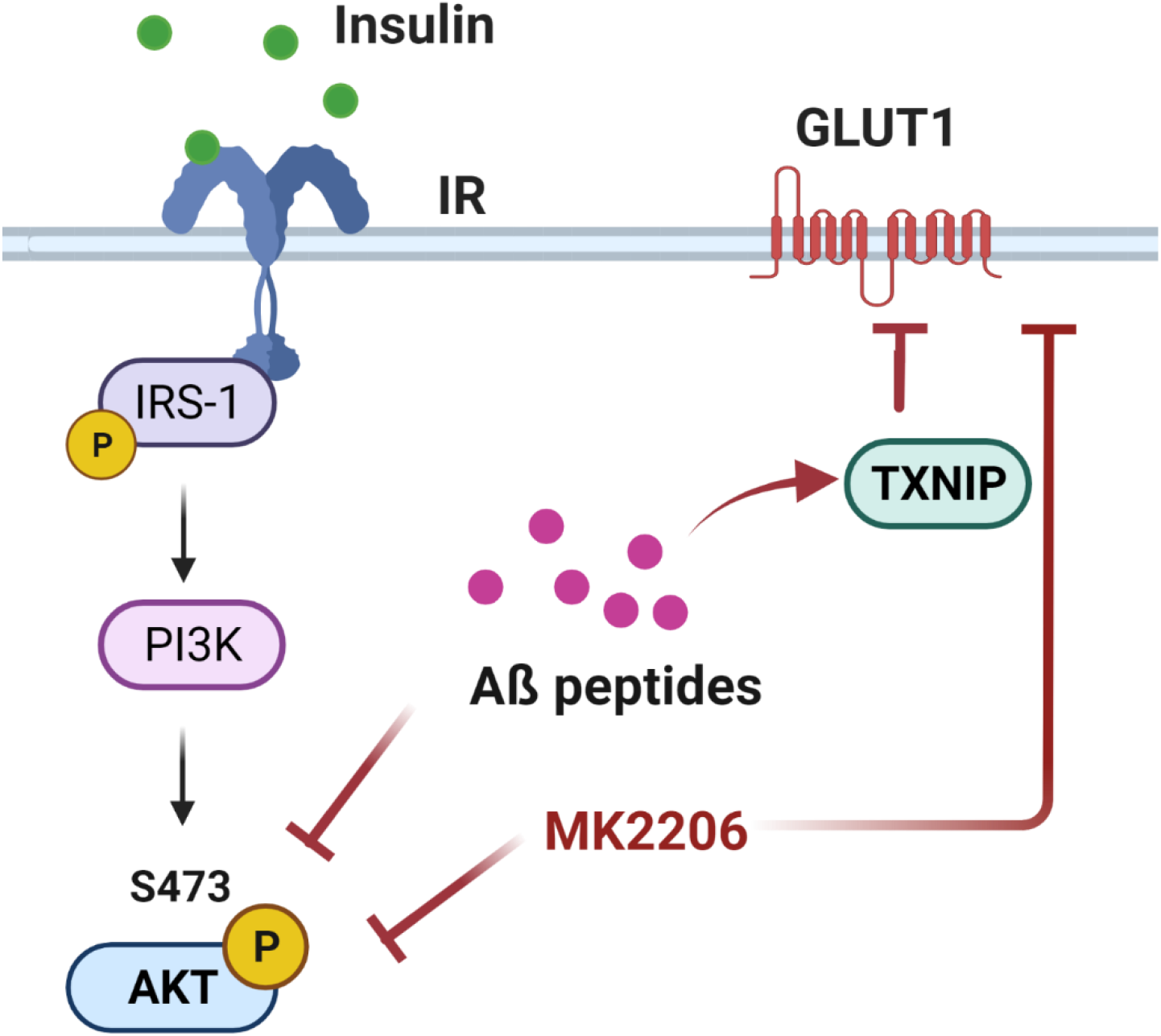
Schematic plot showing sAß peptides suppress insulin-Akt pathway to inhibit GLUT1 expression through TXNIIP.

## Author Contributions

*Participated in research design:* LW, LL, VJL, KKK

*Conducted experiments:* LW, GLC

*Contributed analytic tools:* PHM

*Performed data analysis:* LW

*Wrote or contributed to the writing of the manuscript:* All authors

